# *Azospirillum brasilense* and tomato exudate or cytidine increase phytopathogen resistance and modulate phyllosphere/rhizosphere

**DOI:** 10.1101/2025.10.08.680754

**Authors:** Janice N.A. Tagoe, Bidya Ojha, Shelley M. Horne, Birgit M. Prüß

**Affiliations:** Department of Plant Pathology, Microbiology, and Biotechnology, North Dakota State University, Fargo, ND, 58108

**Keywords:** plant growth beneficial rhizobacteria, *Azospirillum brasilense*, cytidine, tomatoes, *Pseudomonas syringae*, phyllosphere, rhizosphere, human and plant pathogens

## Abstract

Tomatoes are an important crop worldwide and phytopathogens can cause devastating losses. This study describes a treatment, consisting of *Azospirillum brasilense* Sp7 and either tomato seedling exudate or the exudate compound cytidine. The combination of *A. brasilense* Sp7 with cytidine showed a remarkable reduction of 83.4% in disease severity of tomatoes challenged with *Pseudomonas syringae* pv. *tomato* DC3000. Replacing cytidine with exudate was less effective at 71%, but the reduction in disease severity was still larger than by *A. brasilense* Sp7 alone at 55%. This reduction in disease severity was not paralleled by a decrease in *P. syringae* in leaf homogenates. Cytidine caused a 6.7 fold increase in *A. brasilense* Sp7 16S rDNA in root homogenates.

In phyllosphere and rhizosphere, treatments modulated the microbial composition. In the phyllosphere, specific associations between treatment groups and bacterial orders could be computed. In the rhizosphere, principal component analysis revealed that variation along PC1 was dominated by the presence or absence of *A. brasilense*. Intriguingly, the inoculant caused an increase in the abundance of other Azospirillales species.

Les tomates sont une culture d’importance mondiale, et les phytopathogènes peuvent entraîner des pertes dévastatrices. Notre étude décrit un traitement composé *d’Azospirillum brasilense* Sp7 et soit d’un exsudat de plantules de tomate, soit de la cytidine, un composé particulier provenant des exsudats et supposé avoir un effet chimiotactique. L’association de *A. brasilense* Sp7 avec la cytidine a entraîné une réduction remarquable de 83,4 % de la sévérité de la maladie parmi les plantes de tomate infectées par *Pseudomonas syringae* pv. *tomato* DC3000. Le remplacement de la cytidine par des exsudats était moins efficace (71 %), bien que la réduction de la sévérité de la maladie ait demeuré supérieure à celle obtenue avec *A. brasilense* Sp7 seul (55 %). Cette diminution de la sévérité de la maladie n’a toutefois pas été accompagnée d’une réduction de la concentration de *P. syringae* dans les homogénats foliaires. La cytidine a provoqué une augmentation de 6,7 fois de l’ADNr 16S de *A. brasilense* Sp7 dans les homogénats racinaires.

Dans la phyllosphère et la rhizosphère, les traitements ont modulé la composition microbienne. Dans la phyllosphère, des associations spécifiques entre les groupes du traitement et les ordres bactériens ont pu être calculées. Dans la rhizosphère, l’analyse en composantes principales a montré que la variation de l’axe PC1 était dominée par la présence ou l’absence d’*A. brasilense*. De manière intéressante, l’inoculum a entraîné une augmentation de l’abondance d’autres espèces d’Azospirillales.

## Introduction

Tomatoes (*Solanum lycopersicum*) are an economically important vegetable crop worldwide (Bergougnoux 2014); losses occur through phytopathogen infections, both fungal (*e.g. Fusarium oxysporum* f. sp. *lycopersici* (Ma et al. 2013)) or bacterial (*e.g. Pseudomonas syringae* pv. *tomato* (El-Fatah et al. 2023)). The latter has been used as a model organism for understanding phytopathogen infection in tomatoes since the 1980s. *P. syringae* pv. *tomato* DC3000 was isolated in 1991 from *Arabidopsis thaliana* (Whalen et al. 1991).

Protection of tomatoes against phytopathogens can be mediated by natural components of the soil microflora designated <u>P</u>lant <u>G</u>rowth <u>P</u>romoting <u>R</u>hizobacteria (PGPR). This includes other species of *Pseudomonas*, such as *Pseudomonas resinovorans*, *Pseudomonas vranovensis*, or *Pseudomonas putida* (Elsharkawy et al. 2023). *Azospirillum brasilense* is currently recognized as a PGPR with ecological function and agricultural biotechnology innovation (Cassán et al. 2026; Cassán 2026). First identified as forming associations with plants in grasslands (Steenhoudt and Vanderleyden 2000), its uses have extended to monocot crop plants like wheat (*Tritivum aestivum*, (Pereg-Gerk et al. 1998; Vazquez et al. 2021)) or maize (*Zea mays*, (Vidotti et al. 2019)) and dicots including tomatoes (El-Beltagi et al. 2022; Pellegrini et al. 2020). *A. brasilense* colonizes assorted plant tissues, including maize roots (da Cunha et al. 2020) or tomato leaves (Botta et al. 2013). In addition to the nitrogenase system, which is induced under nitrogen limiting conditions (Housh et al. 2023; Wang et al. 2024), benefits from *A. brasilense* colonization for the plants include phytohormone production, changes in root architecture, increased nutrient and water uptake, enhanced leaf development, and increased crop productivity (Cassán et al. 2014; Contreras-Cornejo 2026). Inoculation with *A. brasilense* reshapes the plant microbiome, including a reduction in the abundance of several pathogens (Coniglio et al. 2022).

The rhizospheric lifestyle of *A. brasilense* relies on multiple mechanisms for efficient niche colonization, the first of them being motility and chemotaxis of soil bacteria towards plant exuded metabolites (Nievas et al. 2023). The composition of tomato exudates and chemotaxis towards exudate compounds has been extensively studied (Alexandre 2026; Alexandre et al. 2000; Ganusova et al. 2024; Ganusova et al. 2021; Ganusova et al. 2023). Exudates can enhance plant health and phytopathogen resistance. Examples of this include methyl ferulate preventing tobacco black shank disease (Ma et al. 2025), oxylipins that prevent foliar phytopathogens (Huang et al. 2023), glycoalkaloids to modulate microbial communities and defend against pathogens (Nakayasu et al. 2022; Nakayasu et al. 2021), and acyl sugars acting as insecticides (Korenblum et al. 2020).

Another group of plant exudate compounds that improve plant health are nucleosides (Keren et al. 2024). Exuded by *Helianthemum sessiliflorum,* nucleosides were able to travel in a soil plate assay for quite some distances and permitted the recruitment of a range of plant beneficial bacteria and human pathogens. Our own previous studies were consistent with these observations. Cytidine was one of the compounds that were identified in pea, tomato, and cucumber seedling exudate (Nisha et al. 2024). The nucleoside served as chemoattractant for *A. brasilense* on semi-solid swim plates (Nisha et al. 2024) and enhanced root architecture of one week old pea plants (Nisha et al. 2025).

This study tested the effect of a combination treatment that consisted of *A. brasilense* Sp7 and either tomato seedling exudate or the exudate compound cytidine on phytopathogen resistance of tomatoes, as well as the microbial composition of tomato phyllosphere and rhizosphere. In the first part of the study, the hypothesis was tested that tomato seedling exudate or cytidine might enhance the ability of *A. brasilense* Sp7 to aid the tomato plants in resisting infection with *P. syringae* pv. *tomato* DC3000. The effect of the treatments on the abundance of pathogen on leaves and the PGPR on roots was investigated. The second part of the study looked beyond the tri-partite model of PGPR, phytopathogen, and tomato and investigated the effect of the treatments on phyllosphere and rhizosphere of the tomato plants.

## Materials and Methods

### Bacteria and bacterial growth conditions

Bacteria used for this study were *A. brasilense* Sp7 (Eskew 1977), kindly provided by Dr. Gladys Alexandre (University of Tennessee, Knoxville, TN) and *P. syringae* pv. *tomato* DC3000 (ATCC BAA-871) (Buell et al. 2003). *A. brasilense* Sp7 was cultured in <u>T</u>ryptone <u>Y</u>east extract (TY: tryptone, 5 g/l; yeast extract, 3 g/l; CaCl_2_. 6H_2_O, 1.3 g/l (Beringer 1974)). *P. syringae* pv. *tomato* DC3000 was grown in liquid King’s B medium (20 g/l proteose peptone, 1.5 g/l K_2_HPO_4_, 10 ml/l glycerol, 1.5 g/l MgSO_4_. 7 H_2_O), pH 7,2 ± 0.2) (King et al. 1954), supplemented with 50 µg/ml rifampicin. For agar plates, 15 g/l agar were added.

### Tomato seeds and seed germination

Yellow pear tomato seeds (SKU34924) were from True Leaf Market (Salt Lake City, UT). Seeds were surface-sterilized for 30 s in 95% ethanol and 5 min in 2% bleach, then rinsed 10 times with sterile dH_2_O (dH_2_O). Seeds germinated for 3 days on 0.8% agar in dH_2_O. For experiments with seedling exudate, tomato exudate was produced from 1 g of freshly germinated seeds in 10 ml of dH_2_O (Nisha et al. 2024). Seeds germinated under three separate conditions: dH_2_O, seedling exudate at 1.5 mg of exudate dry weight/ml dH_2_O, or cytidine at 100 mM. Germinated seeds were planted when approximately half an inch of root was visible.

### Plant growth

To permit maximum colonization with our inoculant in the initial days of plant growth (Bashan and De-Bashan 2002; King et al. 2024; Querejeta 2023), growth medium was sterilized by autoclaving. The medium was a 1:1 (w/w) mix of soil (Promix BX General Purpose, 10280; Premier Horticulture Inc. Canada, Québec, Canada) and vermiculite (Vermiculite Medium PVP; PVP Industries, North Bloomfield, OH). Germinated seeds were planted into the center of pots of ∼4 inch in diameter and 6 inch in height. *A. brasilense* Sp7 was inoculated immediately to the location of the planted seed at 10^3^ to 10^4^ CFU in 1 ml of dH_2_O. Plants were grown in a Gen 2000 growth chamber (Conviron, Pembina, ND) at 26°C and light for 16 hours. Tomato-tone (The Espoma Company, Millville, NJ) was used as fertilizer following the manufacturer’s instructions.

### Phytopathogen experiment and disease assessment

The backbone of the phytopathogen experiments are the studies by the laboratories of Dr. Emilia Lopez-Solanilla (Cerna-Vargas et al. 2019), Dr. Melanie Filiatrault (Chakravarthy et al. 2017), and Drs. Yoav Bashan and Luz de-Bashan (Bashan and De-Bashan 2002); the original protocols were modified by the supplementation with cytidine or total seedling exudate in the germination step and *A. brasilense* Sp7 at planting.

Briefly, tomato seedlings germinated in dH_2_O, exudate, or cytidine were planted and inoculated with/without *A. brasilense* Sp7. Four weeks old tomato plants were dip-inoculated with *P. syringae* pv. *tomato* DC3000 by dipping for 30 s in 3 x 10^7^ CFU ml^-1^ in 10 mM MgCl_2_, then air dried and incubated in a Gen 2000 growth chamber at 26°C with 80% relative humidity and 16 h of light. Control groups of un-challenged plants were dipped in MgCl_2_ as part of the first experiment and compared to un-dipped plants. Neither of these plants exhibited disease symptoms and live counts were similar. For this reason, un-challenged control groups in the consecutive experiments were no longer dipped. Six days after the challenge, plants were photographed and assessed for disease severity.

Disease severity data were collected from three replicate experiments in twelve test groups. Symptoms started with dark necrotic lesions on leaves, then leaf wilting. In severe cases, this was followed by leaf and ultimately plant death. Disease severity was expressed as percent death of the leaves from each plant. One healthy plant that had not been challenged with the pathogen and one severely diseased plant were used as comparisons. The assessment was done by at least two researchers in steps of 10%. Examples of plants that were assessed as 0% dead, 50% dead, and 90% dead are presented in Figure S1.

For *Pseudomonas* counts, data were collected from two replicate phytopathogen experiments. Live bacterial counts were obtained from homogenates that were produced from 6 leaf discs per plant that were punched with a 1/4 inch sterile cork borer. Weighed leaf discs were homogenized in 10 mM MgCl_2_ with pestles. The leaf homogenates were serially diluted and plated onto King’s B agar to determine *Pseudomonas* counts (King et al. 1954). Bacterial counts were expressed as log_10_ CFU/g of plant material.

For disease severity and *Pseudomonas* counts, data were presented as Box and Whiskers generated with GraphPad Prism 11.0.0 (84). The lines within the boxes indicate the medians of the populations. Data points from individual plants are included, means of the populations are given for pathogen challenged test groups. Statistical analysis was done by one-way ANOVA, followed with Tukey for pairwise comparisons between test groups. A *p*-value below 0.05 was indicative of a statistically significant difference between test groups. Percent disease reduction was calculated as 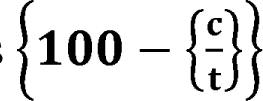, where t are the mean percent death data from the treatment groups and c the mean percent death data from the untreated control group (-*Abra* +Ps).

### 16S rDNA analysis

For the determination of bacterial communities, gDNA was isolated from leaf and root homogenates from three plants from one phytopathogen experiment. Prior to homogenizing, the roots were washed with dH_2_O thoroughly but carefully, so that most of the soil would be gone, but the outside of the roots would still be intact. This sampling would yield the epiphytic and endophytic microbiomes of the roots.

Homogenates were produced from approximately 50 mg of plant material by using the Cole-Parmer HG-200 GenoLyteR Compact Homogenizer (Cole-Parmer, Vernon Hills, IL) at 2,000 rpm for 1 min. DNA extraction was done with the Qiagen Plant Pro kit (Qiagen, Hilden, Germany). Quality of the DNA samples was analyzed with the Nanodrop One Spectrophotometer, quantity with the Qubit 4 Fluorometer (Fisher Scientific, Hampton, NH). Samples were diluted to 1 ng/µl. The construction of the library and 16S rDNA sequencing were done at the Dr. Thomas Glass Biotech Innovation Core at NDSU. The QIAseq 16S/ITS Region Panel (Qiagen) was used for the generation of the 16S rDNA library. The primers used were 341f:CCTACGGGNGGCWGCAG and 785r:GACTACHVGGGTATCTAATCC, which amplify the V3/V4 hyper variable region of the 16S rDNA (Qiagen). The PCR protocol from the manufacturer was followed and PCR products were cleaned with QIAseq bead cleanup. The index PCR reaction was done with unique dual indexes for each sample, PCR products were cleaned with QIAseq bead cleanup. Samples were quantified with the Qubit 4 Fluorometer and analyzed with the 4200 Tape Station (Agilent, Santa Clara, CA). Sequencing was performed using the P1 XLEAP-SBS 600 cycle sequencing kit on the NextSeq 1000 (Illumina, San Diego, CA).

Phyllosphere and rhizosphere data were analyzed separately. Fastq files were imported into CLC Genomics Workbench (Qiagen) for analysis using the Microbial Genomics Module. The Data QC and OTV clustering workflow was used to trim adaptors, poly G regions, ambiguities and sequences not meeting the quality threshold of 0.05. Reference based OTV clustering using the Silva SSU 99% (138.2) database was performed using the similarity percent specified by the database. Minimum occurrence was set at 2. The species identifiers or taxonomies in the CLC Workbench are from the Earth Microbiome Project (https://earthmicrobiome.org/). The resulting abundance tables were exported to Excel separately for phyllosphere and rhizosphere and further analyzed. These two tables are provided as online supplements and include the ASV (Tables S1 and S2). Rarefaction curves for both data sets are shown in Figure S2.

Before analyzing the data, the taxonomies from the inoculant *A. brasilense* Sp7 and the challenge *P. syringae* pv. *tomato* DC3000 needed to be confirmed/identified. Liquid pure cultures of each strain were grown and 16S rDNA was sequenced. This led to the conclusion that the inoculant *A. brasilense* Sp7 was *A. brasilense* AF533354.1.1439 and the challenge *P. syringae* pv. *tomato* DC3000 was *P. amygdali* pv. *morsprunorum* AB0011445.1.1538. As a final pre-analysis step, small abundance taxonomies were filtered out. In the phyllosphere, ≤49 across all test groups and replicates was used as threshold to eliminate such small abundances. In the rhizosphere, this threshold was ≤499. Plant and fungal taxonomies were eliminated.

As part of the phytopathogen experiment, 16S rDNA abundances from the challenge pathogen *P. syringae* pv. *tomato* DC3000 in the phyllosphere and the inoculant *A. brasilense* Sp7 in the rhizosphere were analyzed. The data sets were statistically analyzed with one-way ANOVA to compare either all 12 test groups or just the six pathogen challenged test groups (+Ps). Data were presented as floating bars in GraphPad Prism.

Analysis of the phyllosphere was started with the generation of a phylogenetic tree for 67 phyllosphere taxonomies. ASV were converted into a FASTA file and loaded into Galaxy (usegalaxy.eu/). Multiple sequence alignment was done with MAFFT, followed by FASTTREE. The Newick file (.nhx) from the maximum likelihood computation was used in iTOL (itol.embl.de). Using the mid-point root, the circular tree was selected.

The next step was a comprehensive analysis of 16S rDNA abundances within phyllosphere and rhizosphere. First, richness was computed, separately for each replicate in Excel with the following function: =countif(A2:A69, “>1”). Second, relative abundances were summed over the 67 taxonomies from the phyllosphere and the 280 taxonomies from the rhizosphere, also separately for each of the three replicates in Excel. For both analyses, data were transferred into GraphPad Prism and graphed as floating bars. Third, for the generation of the heatmaps, 16S rDNA abundances from taxonomies that represent the different bacterial orders were summed across all species. Mean percent abundances across the three replicates were used and visualized in GraphPad Prism. For the phyllosphere, only pathogen challenged test groups were included in the heatmap. Fourth, the same data set was used for principal component analysis (PCA) in GraphPad Prism, graphing PC scores (treatments) and Biplot analysis (treatment and orders). Association strength was computed between treatments and orders with cosinus similarity.

### Digital PCR

From the same DNA samples that were used for 16S rDNA sequencing, digital PCR was performed to confirm 16S rDNA data for our challenge pathogen *P. syringae* pv. *Tomato* DC3000. Primers for *P. syringae* pv. *tomato* DC3000 were PST-*hrpL*_e_fwd, 5’-TTTCAACATGCCAGCAAACC-3’ and PST-*hrpL*_e_rev, 5’-GATGCCCCTCTACCTGATGA-3’ as described (Peňázová et al. 2020). The probe was PST-*hrpL*_TP, 5’-GCTGAACCTGATCCGCAATCAC-3’, marked with the yellow fluorescence dye HEX.

Each PCR reaction mix of 40 µl contained between 50 and 200 ng of DNA, 1 x PCR mix, 0.8 µM primer mix, 0.4 µM probe, and 0.25 U EcoRI in RNAse free water. The PCR reactions were performed in a QIAcuity Digital PCR System (Qiagen) on 24 well QIAcuity Nanoplates at 40 cycles of annealing/extension at 55°C for 30 s and denaturation at 95°C for 15 s. After 40 cycles, fluorescence was measured in each of the 25,000 partitions of a well. Individual reactions were considered positive when a threshold of 50 to 150 was achieved. The original data were expressed as count of positive and negative reactions in the partitions and computed to copies/µl in the QIAcuity Software Suite v2.1.7.182. Given the start concentration of DNA, these data were converted to copies/ng of DNA. Data were graphed as floating bars in GraphPad Prism, statistical analysis was done with ANOVA.

## Results

### Treatments decreased disease severity on pathogen challenged tomatoes

The first goal of the study was to investigate the effect of the treatments on phytopathogen resistance of the tomato plants. The experiment was done in 12 test groups, covering absence or presence of each of the following: *A. brasilense* Sp7 (*Abra*), *P. syringae* pv. *tomato* DC3000 (Ps), seedling exudate (Ex), or cytidine (CYD). Fig. 1 shows disease severity data expressed in percent death at 6 days post challenge.

**Fig. 1.**
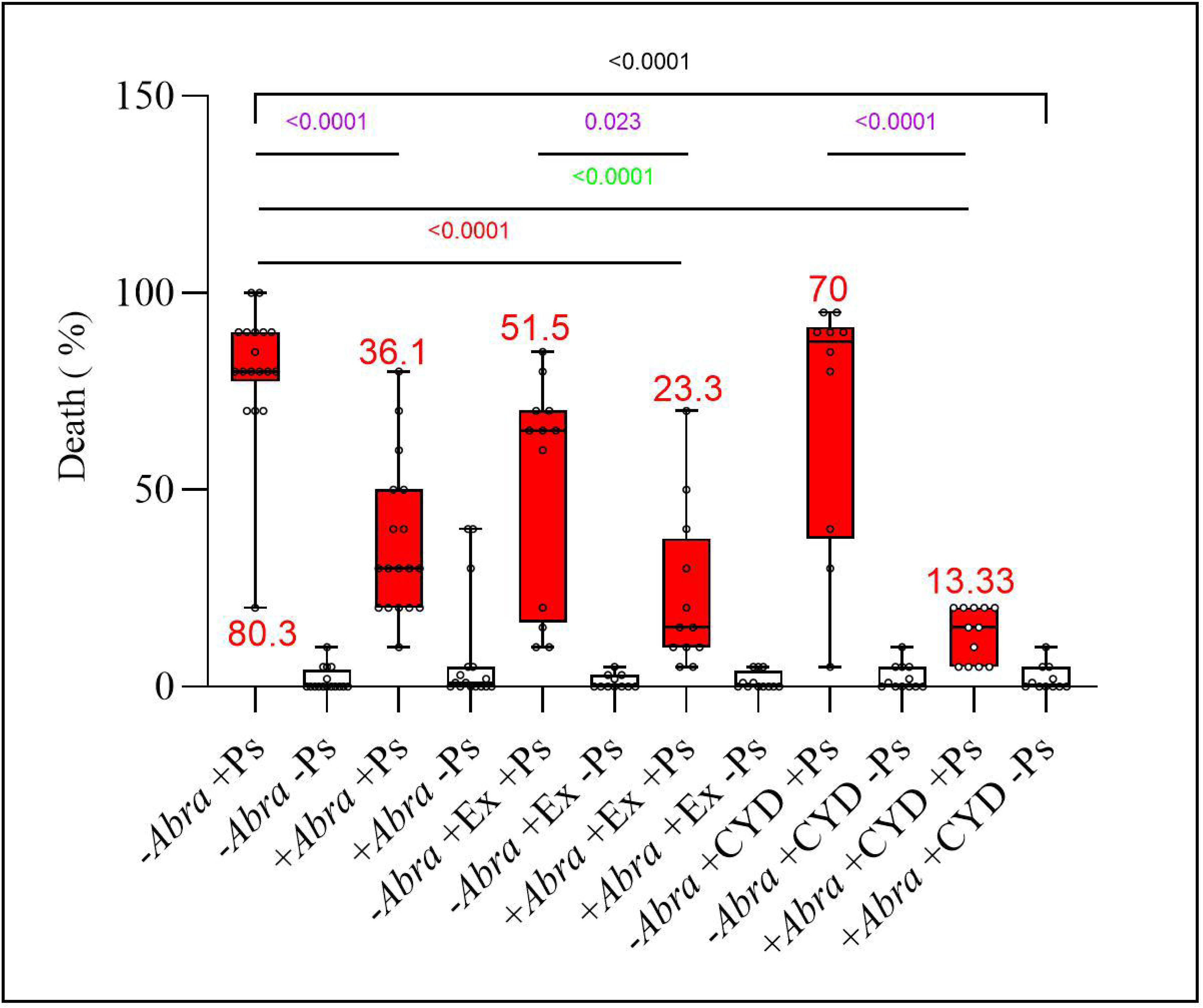
Disease severity. For the disease assessments, percent leaf death data were collected from a minimum of 3 replicate experiments. Red boxes mark data populations for test groups that were challenged with *P. syringae* pv. *tomato* DC3000, white boxes mark data populations for un-challenged test groups. The lines within the boxes indicate the medians of the populations, the small open circles within the boxes are the data points from each individual plant. The numbers in red are the averages across the populations for challenged test groups. The blunt end line across all test groups indicates the *p*-value from the one-way ANOVA. The open end lines between two bars are the *p*-values from pairwise comparisons with the Tukey test.

The *p*-value from the one-way ANOVA across the 12 test groups was <0.0001. These were also the *p*-values from the Tukey pairwise comparisons for the six comparisons between plants challenged or not challenged with *P. syringae* pv. *tomato* DC3000 (*p*-values not included in Fig. 1). In the absence of the pathogen, disease severity ranged between 1% and 2% death (white bars). For the three comparisons between *P. syringae* challenged plants that were either inoculated or not inoculated with *A. brasilense* Sp7 (-*Abra* +Ps versus +*Abra* +Ps; -*Abra* +Ex +Ps versus +*Abra* +Ex +Ps; -*Abra* +CYD +Ps versus +*Abra* +CYD +Ps) (*p*-values in purple), the *p*-values indicated statistical significance of the differences. *A. brasilense* Sp7 had a positive effect on the health of the challenged plants. The -*Abra* +Ps vs +*Abra* +Ps comparison yielded a disease reduction of 55% from 80.3% to 36.1% leaf death.

An improvement in disease reduction by *A. brasilense* Sp7 was accomplished by germinating seeds in seedling exudate (*p*-values in red) and more so in cytidine (*p*-values in green). Comparing the test group -*Abra* +Ps at 80.3% death to +*Abra* +Ex +Ps at 23.3% death, the reduction in disease severity was 71% at a *p*-value indicating statistical significance of the difference. Comparing the test group -*Abra* +Ps at 80.3% death to the test group +*Abra* +Cyd +Ps at 13.3% death, the reduction in disease severity was 83.4% at a *p*-value below 0.05.

### The treatments did not decrease abundance of the pathogen on leaves, cytidine increased abundance of the PGPR on roots

The reduction in disease severity by the treatments could have multiple causes; among the possibilities are a decrease in the abundance of *P. syringae* on leaves or an increase in the abundance of *A. brasilense* on roots.

The abundance of *P. syringae* was determined as *Pseudomonas* counts on rifampicin supplemented King’s B medium, abundance of 16S rDNA by means of sequencing, and copies of the *hrpL* gene by means of digital PCR. Note that King’s B medium detects multiple microorganisms in addition to species from the *Pseudomonas* genus, in particular those that produce fluorescent siderophores (Johnsen and Nielsen 1999), the *hrpL* gene is specific for *P. syringae* (Peňázová et al. 2020), and 16S rDNA is specific for our *P. syringae* pv. *tomato* DC3000.

Fig. 2A shows the log_10_ CFU/g data from the colony counts. Pathogen challenged plants contained a mean of 8.1 to 9.25 log_10_ CFU of *Pseudomonas*/g of plant material (red bars). Un-challenged plants contained a smaller but still remarkable number of *Pseudomonas* and/or other fluorescent bacteria between 4.2 and 7.25 log_10_ CFU/g (white bars). The only statistically significant differences from the pairwise comparisons were five out of six comparisons where a test group of plants challenged with *P. syringae* pv. *tomato* DC3000 was compared to an un-challenged group (*e.g*. -*Abra* +Ps versus -*Abra* -Ps). A second ANOVA was performed to compare the six test groups of challenged plants. The *p*-value for this comparison did not indicate statistically significant differences between challenged test groups.

**Fig. 2.**
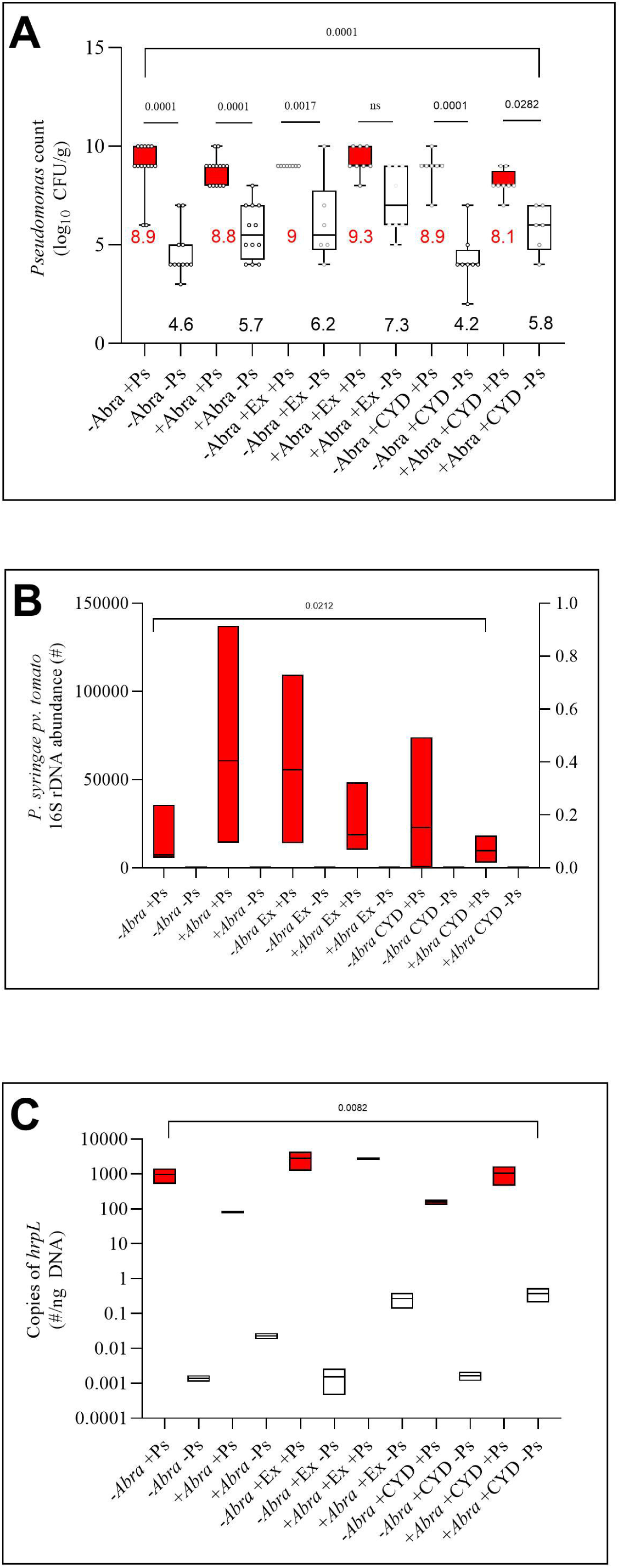
Abundance of *P. syringae* on tomato leaves. Panel A shows the live *Pseudomonas* counts on tomato leaves. Abundance data were collected as log_10_ CFU/g plant material from two phytopathogen experiments. The numbers in red are the averages across the populations for challenged test groups, numbers in black are for un-challenged test groups. Panel B shows the 16S rDNA abundance of *P. syringae* pv. *tomato* DC3000. 16S rDNA was sequenced from leaf homogenates from 3 plants. Panel C shows the number of copies of the *hrpL* gene from the same leaf homogenates by dPCR. For all panels, red boxes mark data for test groups that were challenged with the pathogen, white boxes mark data populations for un-challenged test groups. The lines within the boxes indicate the medians of the populations, the small open circles within the boxes are the data points from each individual plant. The blunt end lines across all test groups indicate the *p*-value from the one-way ANOVA that was performed across all twelve test groups. Open end lines between two bars are *p*-values from pairwise comparisons with Tukey.

Fig. 2B shows data from the 16S rDNA analysis. The *p*-value from the one-way ANOVA that was performed across all twelve test groups indicated statistically significant differences between test groups. However, calculating an ANOVA across only the six un-challenged test groups was 0.1740. The only statistically significant differences were between test groups whose sole treatment difference was presence or absence of the pathogen. Leaf homogenates from un-challenged tomato plants did not contain 16S rDNA from the challenge pathogen.

Fig. 2C shows copies of the *hrpL* gene larger than 1/ng of DNA only for test groups that had been challenged with the pathogen (red bars). The ANOVA that was computed across all 12 test groups did point towards statistically significant differences with a *p*-value of 0.0082. The ANOVA across the six pathogen challenged test groups was 0.1157. In three experiments, the treatments did not reduce the abundance of *P. syringae*.

For the abundance of *A. brasilense* on roots, the same types of experiments were performed as for the abundance of *P. syringae* from root homogenates, live counts from TY strep plates, 16S rDNA, and dPCR (Reddy Priya et al. 2016). Fig. 3 shows the 16S rDNA analysis for *A. brasilense* Sp7. The ANOVA that was performed on the complete data set yielded a *p*-value of statistical significance. When comparing the +*Abra* -Ps test group with the +*Abra* +CYD -Ps group, cytidine increased 16S rDNA abundance for *A. brasilense* Sp7 by 6.7 fold with *p*-values of 0.0004 and 0.05 from the parametric *t*-test and the non-parametric Mann-Whitney U test, respectively. TY strep plates did not permit the growth of colonies from the homogenates. dPCR did not yield PCR products.

**Fig. 3.**
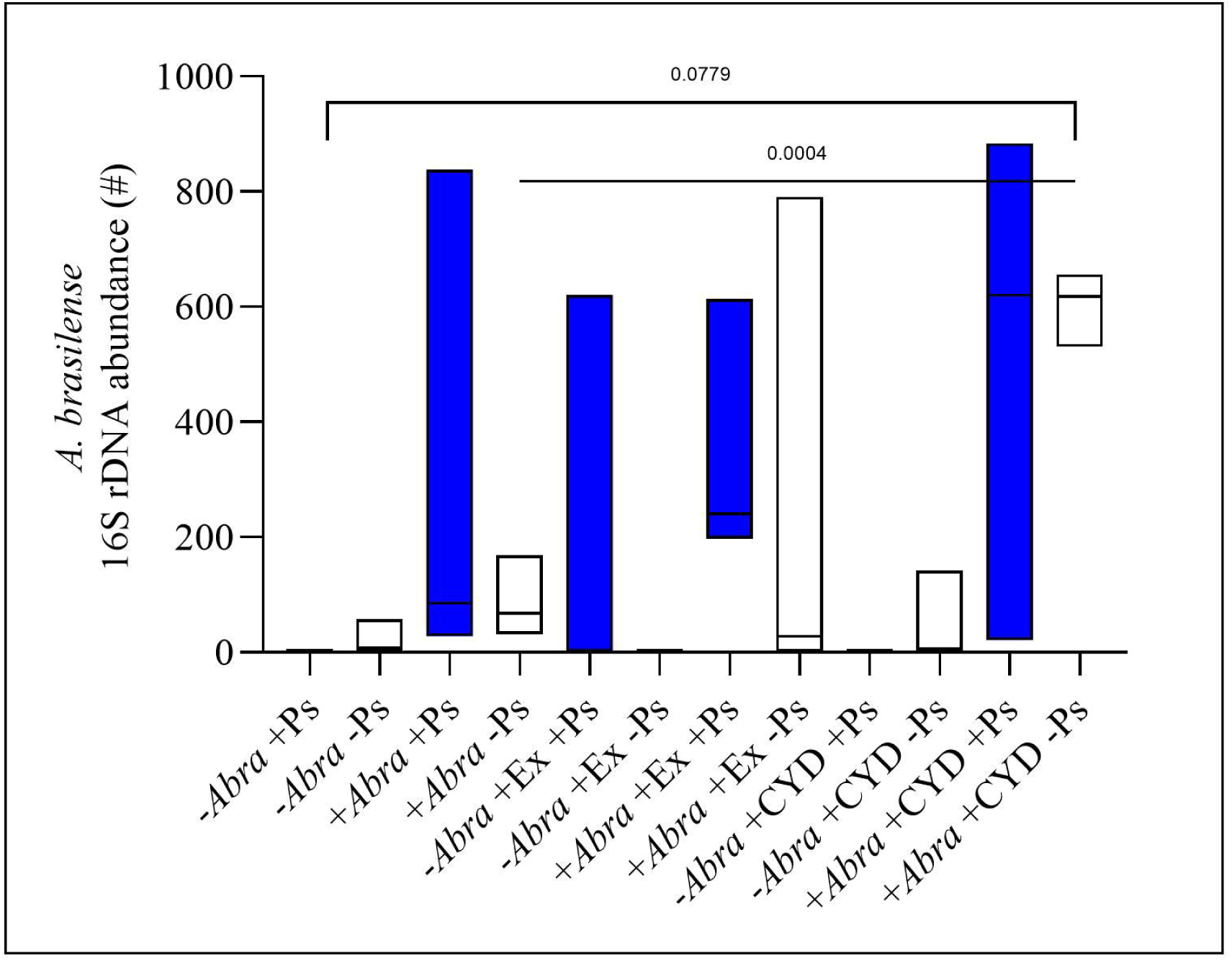
Abundance of *A. brasilense* Sp7 on roots. 16S rDNA was sequenced from root homogenates from 3 plants. Floating bars in blue are from pathogen challenged test groups, floating bars in white are from un-challenged test groups. The line within the floating bars is the median of the population. The blunt end line across all test groups indicates the *p*-value from the one-way ANOVA that was performed across all twelve test groups. The open ended line between two bars is the *p*-value from pairwise comparison with Tukey.

### Treatments modulated the phyllosphere of phytopathogen challenged tomatoes

The second goal of this study was to determine the effect of the treatments on the tomato microbiome. The first analysis was designed to get a glance at the microbial diversity included in the tomato microbiome across all treatment groups to permit comparison with published data (Dong et al. 2019; Romero et al. 2014).

The phyllosphere (Table S1) initially contained 685 taxonomies, which was narrowed down to 67, applying the >50 filter. Fig. 4 shows the phylogenetic tree including these 67 taxonomies that were represented in any of the 12 test groups. The taxonomies clustered into two phyla, Proteobacteria and Bacteroidota. There are 5 species of Sphingomonadales (purple) and 5 species of Rhizobiales (green) among the Alphaproteobacteria, 3 species of Burkholderiales (brown) from the Betaproteobacteria, as well as 2 species of Xanthomonadales (dark purple), 24 species of Pseudomonadales (blue), and 28 species of Enterobacteria (orange) among the Gammaproteobacteria. The order Enterobacterales included the families of *Erwiniaceae* and *Enterobacteriaceae*, the Pseudomonadales were all from the family *Pseudomonadaceae* and the genus *Pseudomonas*. The two species of Bacteroidota were *Flavobacterium ginsengiterrae* from the order Flavobacteriales (yellow) and *Pedobacter kyungeensis* from the order Sphingobacteriales (red). Overall, leaf homogenates included predominantly species from the Proteobacteria.

**Fig. 4.**
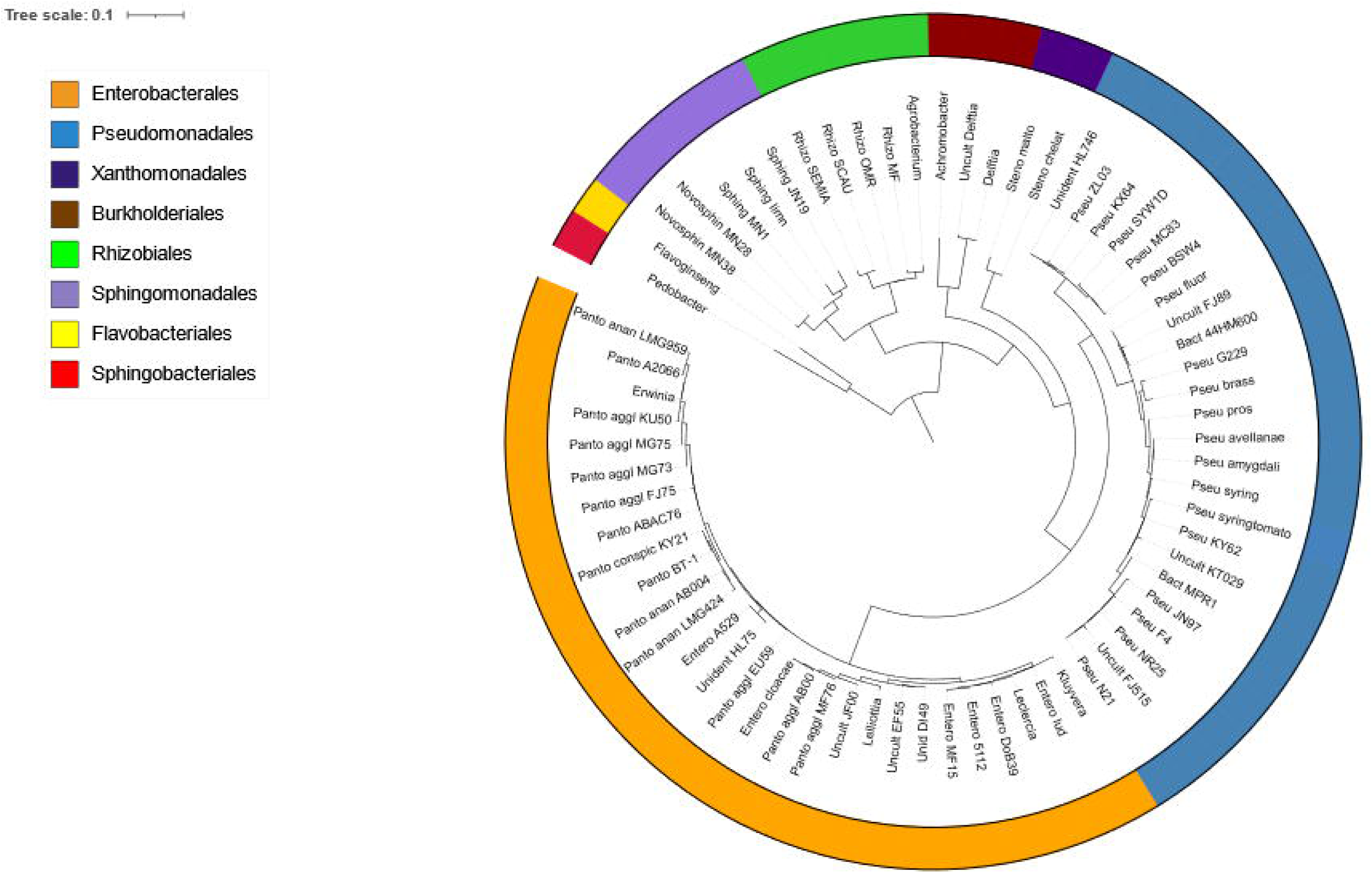
Phylogenetic tree from the tomato phyllosphere. 16S rDNA was analyzed from leaf homogenates from 3 plants. The phylogenetic tree was generated in Galaxy and visualized in iTOL from the 67 bacterial species that exhibited 16S rDNA abundances higher than 50 across all test groups and replicates.

The following analyses were targeted towards understanding the effect of the treatments on microbial composition. Richness (Fig. 5A) and sum abundances (Fig. 5B) were computed separately for each of the 3 replicates/test groups from the 67 phyllosphere taxonomies. Both analyses pointed towards an almost complete lack of 16S rDNA within the phyllosphere of tomatoes that had not been challenged with *P. syringae* pv. *tomato* DC3000. Plants from the test groups -*Abra* +Ex -Ps and -*Abr*a +CYD -Ps harbored a small number of species (Fig. 5A). For richness (Fig. 5A), five out of six comparisons between test groups whose only treatment difference was challenged or un-challenged with the pathogen had a *p*-value below 0.05. For sum abundances (Fig. 5B), the *p*-value from the one-way ANOVA indicated that differences between test groups were lacking statistical significance. By all appearances, the presence of the pathogen permits rise to a myriad of bacteria. Among the pathogen challenged test groups, +*Abra* +Ex +Ps exhibited low richness and low sum abundance. The test groups -*Abra* + Ps and +*Abra* +CYD +Ps had low sum abundances (Fig. 5B).

**Fig. 5.**
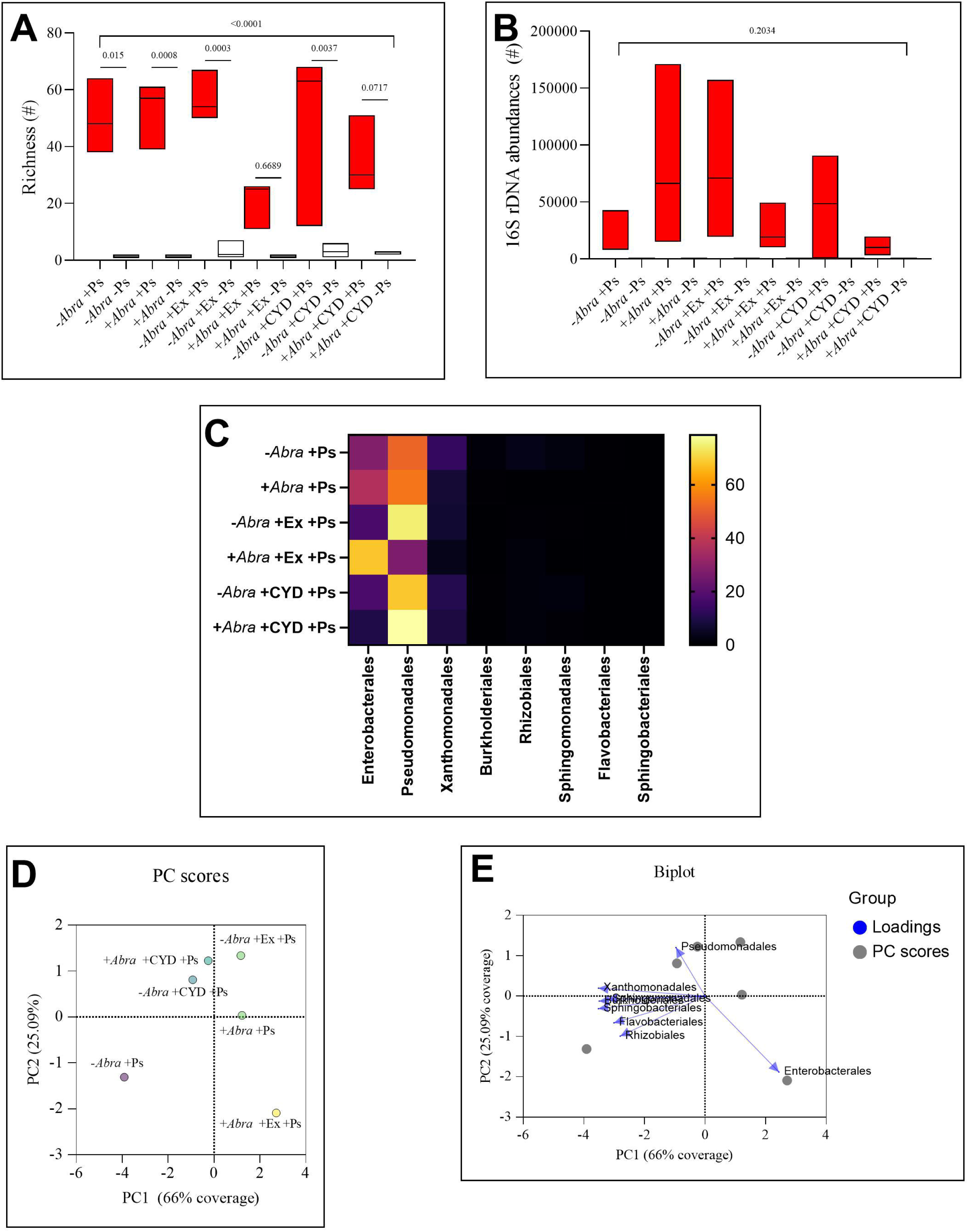
Tomato phyllosphere. Fig. 5A demonstrates richness, Fig. 5B the sum abundances across all bacterial species per test group. Fig. 5C is the heat map for mean percent 16S rDNA abundances from all species within eight orders for pathogen challenged test groups. PCA analysis is presented as PC scores in Fig. 5D and Biplot in Fig. 5E.

Fig. 5C visualizes the percentage of 16S rDNA abundances that were summed across all taxonomies from each of the 8 orders and expressed as percent. Enterobacterales, Pseudomonadales, and Xanthomonadales were the only ones that had significant relative abundances. The highest abundance for Enterobacterales was seen in the +*Abra* +Ex +Ps test group at 68%. There was a reduction in the abundance of 16S rDNA in the +*Abra* +CYD +Ps test group at 9.8% relative to the -*Abra* +CYD +Ps group at 17.4%. For Pseudomonadales, there was a reduction in abundance in the +*Abra* +Ex +Ps test group at 27% relative to the -*Abra* +Ex +Ps group at 75%. For Xanthomonadales, there was a reduction in abundance in all three +*Abra* test groups relative to the respective -*Abra* group.

To investigate associations between test groups and abundances of bacterial orders, PCA was performed, using the mean of the percent abundances (Figs. 5D and E). PC1 captured 66% of the total variation and PC2 captured an additional 25.09% to a cumulative coverage of 91.08%. The six treatments grouped distinctly within the PC space, implying strong multivariate separation between treatments (Fig. 5D). The furthest outlier appeared to be the +*Abra* +Ex +Ps group (yellow) with the PC1 coordinate 2.71 and the PC2 coordinate -2.091. Enterobacterales associated strongly with this +*Abra* +Ex +Ps test group (yellow) in the biplot (Fig. 5E) with a cosinus similarity of 0.924. Pseudomonadales associated with the +*Abra* +CYD +Ps group with a cosinus similarity of 0.998 and with the -*Abra* +CYD +PS group with a cosinus similarity of 0.836 (both light blue in Fig. 5D and E). Across the five analyses that are covered with Fig. 5, the +*Abra* +Ex +Ps and +*Abra* +CYD +Ps test groups stood out, indicating that the respective treatments modulate the phyllosphere.

### Treatments modulated the rhizosphere of phytopathogen challenged tomatoes

In similarity to the phyllosphere, the first rhizosphere analysis was designed to get a glance of the microbial diversity included in the tomato microbiome across all treatment groups to permit comparison with published data. An initial 8,619 taxonomies were narrowed down to 280, applying the much more restrictive >500 filter (Table S2). Among the proteobacteria, there were 93 species of Alphaproteobacteria, 42 species of Betaproteobacteria, and 43 species of Gammaproteobacteria. The non-proteobacteria clustered into eight phyla; Bacteroidota with 61 species, Actinobacteriota with 16 species, Bdellovibrionata with 10 species, Planctomycetota with 4 species, Patescibacteria with 3 species, Myxococcota with 2 species, and Abditibacteriota and Acidobacteria with 1 species each.

To determine the effects of the treatments on microbial composition, richness (Fig. 6A) and 16S rDNA abundances (Fig. 6B) were computed from all 280 taxonomies within the rhizosphere. Both, richness (Fig. 6A) and sum abundances (Fig. 6B) showed 16S rDNA within the rhizosphere of all twelve test groups of plants. For both analyses, the *p*-value from the one-way ANOVAs pointed towards a lack of statistical significance of any differences.

**Fig. 6.**
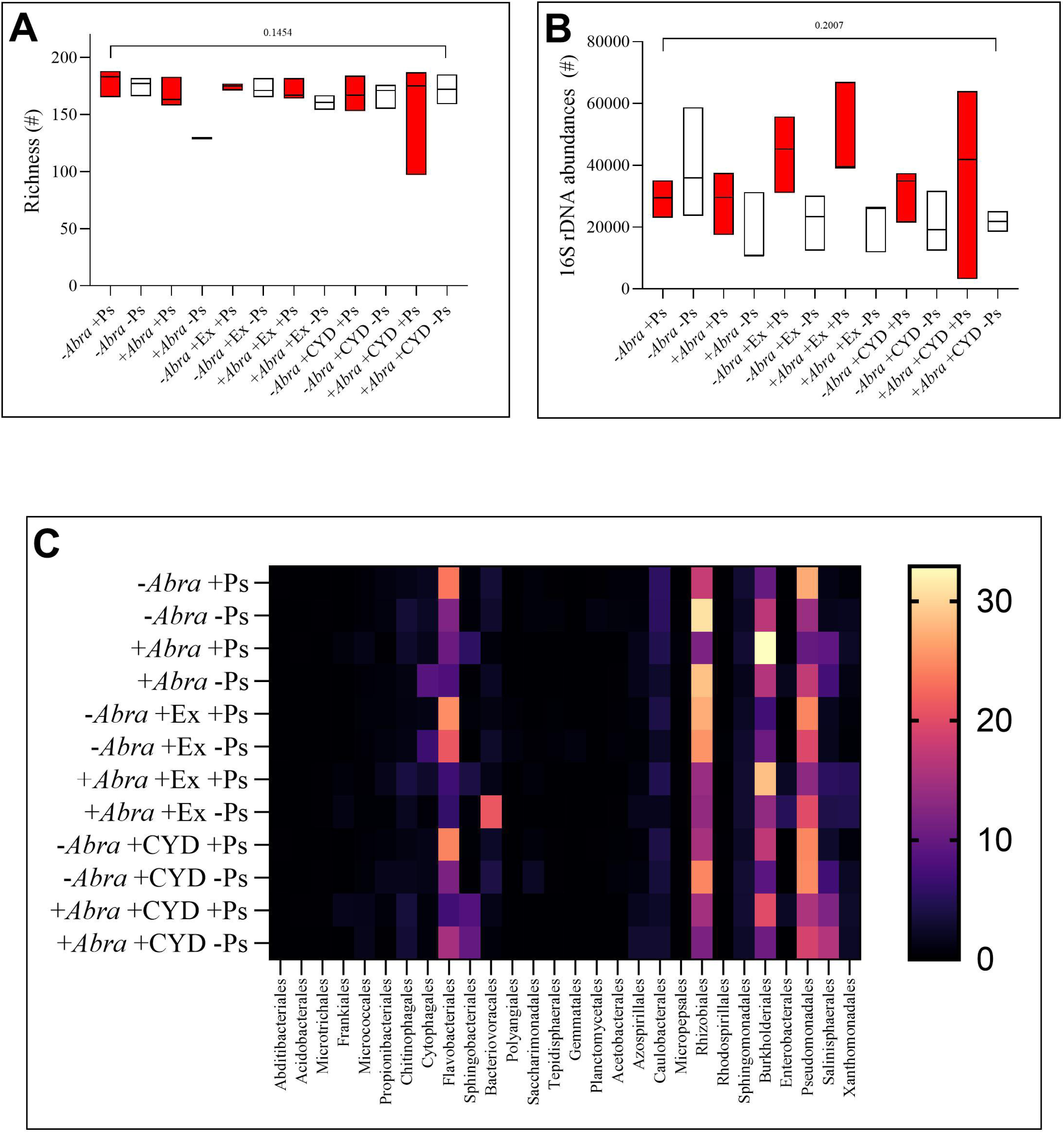
Tomato rhizosphere, richness and abundances. Fig. 6A demonstrates richness, Fig. 6B the sum abundances across all bacterial species per test group. Fig. 6C is the heat map for mean percent 16rDNA abundances from all species within each order.

Fig. 6C shows the heatmap for the 29 bacterial orders in percentage of 16S rDNA abundances. One highly abundant order were the Rhizobiales, especially in the -*Abra* -Ps group at 30.85%, in the +*Abra* -Ps group at 28.59%, the -*Abra* +Ex +Ps group at 27.27%, the -*Abra* +Ex -Ps group at 25.52%, and the -*Abra* +CYD -PS group at 24.5%. Pseudomonadales were high (>24%) in the -*Abra* +Ps, -*Abra* +Ex +Ps, -*Abra* +CYD +Ps and -*Abra* +CYD -Ps groups and lower in the +*Abra* groups. Burkholderiales had the highest percent abundance among any bacterial orders in the +*Abra* +Ps group at 32.9%. In the +*Abra* +Ex +Ps group, Burkholderiales were still high at 28.46%. Flavobacteriales were high in the -*Abra* -Ps group at 23.55%, the - *Abra* +Ex +Ps group at 25.05%, and the -*Abra* +CYD +Ps group at 24.15%. Bacteriovoracales were high in the -*Abra* +Ex –Ps group at 21.04%.

Fig. 7 shows the results from the PCA analysis that was computed from the means of the percent abundances. In Figs. 7A and 7B, PC1 captured 35.65% of the total variation and PC2 captured an additional 15.05% to a cumulative coverage of 50.7%. Treatments with “+A” resembling the presence of *A. brasilense* Sp7 were located on the positive side of the y-axis, while treatments designated “-A” resembling the absence of the inoculant clustered on the negative side (Fig. 7A). This suggests that PC1 was strongly influenced by presence or absence of the PGPR. From the PCA scores, association strengths were calculated as cosinus similarity. Micrococcales, Azospirillales, Salinisphaerales, and Microtrichales associated positively with the presence of *A. brasilense* Sp7 (Table S4). Flavobacteriales, Pseudomonadales, Propionibacteriales, Saccharomonadales, Rhizobiales, Tepidisphaerales, and Abditibacterales associated positively with the absence of *A. brasilense* Sp7 (Table S4).

**Fig. 7.**
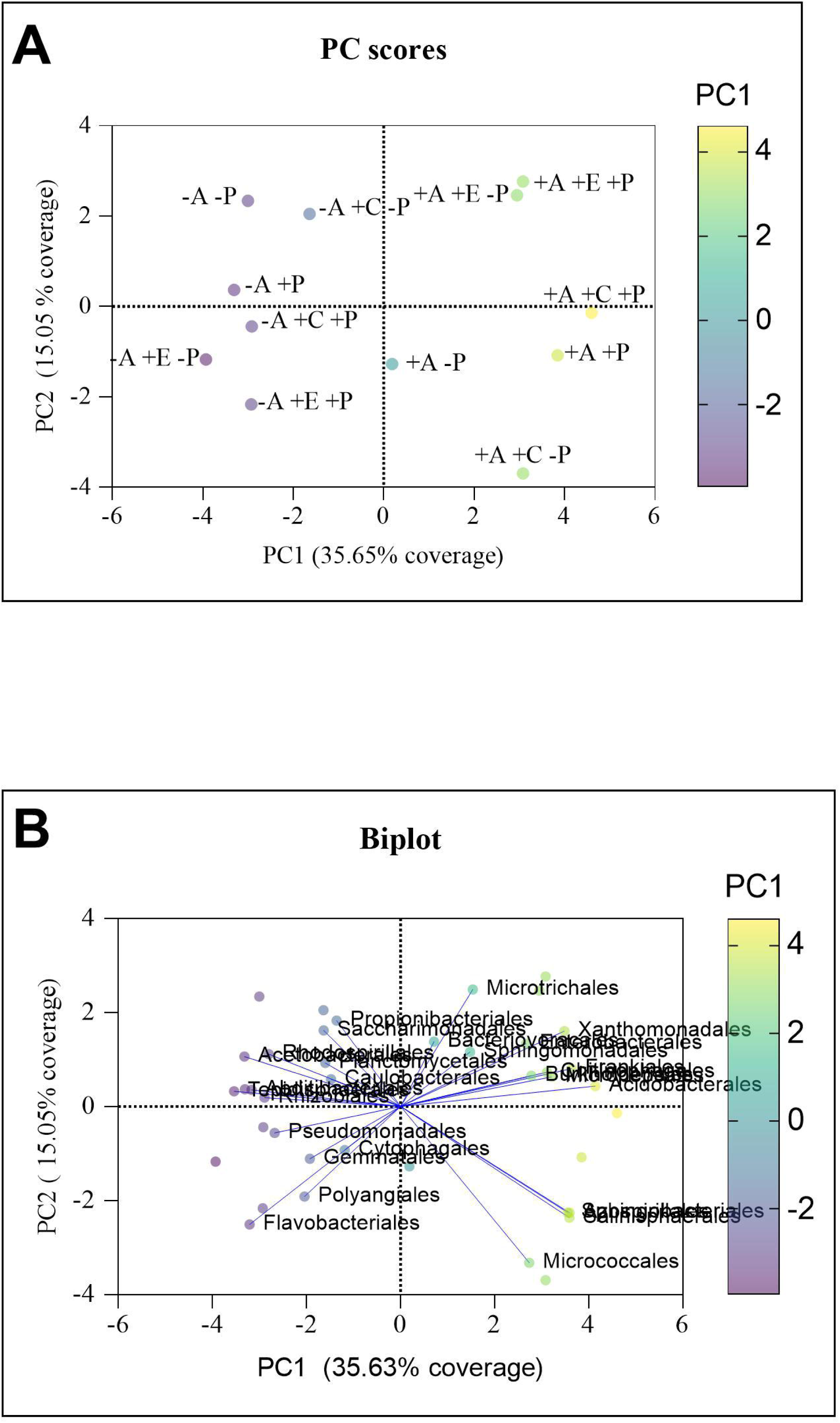
PCA analysis. Fig. 7A shows the PC scores and Fig. 7B the Biplot. See Table S4 for association strengths.

## Discussion

The combination treatments of *A. brasilense* Sp7 and tomato seedling exudate or cytidine reduced disease severity of tomatoes after challenge with *P. syringae* pv. *tomato* DC3000. This effect was larger for the treatment that included cytidine than the one including total exudate.

The reduction in disease severity was not paralleled by a decrease in the abundance of *P. syringae* on leaves, but by an increase in *A. brasilense* Sp7 on roots by cytidine. The treatments modulated the relative abundances of several bacterial orders within phyllosphere and rhizosphere.

## Phytopathogen resistance

Tomatoes are capable of exuding a wide array of metabolic intermediates into the soil to promote their own health and disease resistance, a recent review article summarized exudates as a bridge between roots and rhizospheric microbes (Feng et al. 2024). The general observation that the treatments suppressed disease severity is consistent with many previous studies on disease suppression by PGPR and exudate compounds (Chakravarthy et al. 2017; Janda et al. 2023; Lindeberg et al. 2009; Namgung et al. 2019; Wei and Collmer 2018; Xin and He 2013). In comparison to previous studies, the disease reduction in this study is impressive at 83.4%. Possible causes for this high reduction are numerous. One obvious cause could be a decrease in the abundance of pathogen on the leaves, as was seen for inoculation of tomatoes with *A. brasilense* but without seedling exudate or cytidine (Bashan and De-Bashan 2002). In three different experiments in our study, it was concluded that abundance of the pathogen was not decreased. It is noteworthy that live *Pseudomonas* counts were still high in plants that had not been challenged with the pathogen at 4.2 to 7.2 log_10_ CFU/g. This is because King’s B medium does not just detect *P. syringae*, but other species from the genus *Pseudomonas* and other microorganisms that produce fluorescent siderophores (Johnsen and Nielsen 1999). The 16S rDNA analysis and dPCR (Peňázová et al. 2020) were more selective for *P. syringae* and led to the conclusion that there was little or no *P. syringae* on leaves of plants not challenged with the pathogen.

A second possible cause for disease suppression might be an enhancement of colonization by the inoculant *A. brasilense* Sp7 in response to seedling exudate or cytidine. The phyllosphere did not contain any detectable abundances of *A. brasilense* Sp7 16S rDNA, but cytidine increased *A. brasilense* Sp7 16S rDNA in the rhizosphere by 6.7 fold. This increase was only seen in the 16S rDNA experiment, as live bacterial counts and dPCR failed to detect signals for *A. brasilense* Sp7. This may be due to low abundances of *A. brasilense*, a problem that has been observed before (Compant et al. 2025).

An increase in colonization by the PGPR might lead to an increase in the induced systemic response of the tomatoes, a state of enhanced defensive capacity of the plants after colonized by certain PGPR. For *Azospirillum*, a system of fine-tuned hormonal cross-talk also involving flagellin has been proposed to balance immunity with plant growth (Elías et al. 2022; Fukami et al. 2018; Mora et al. 2023). This means an increase in immunity does not necessarily cause a trade-off in decreased plant growth, which is important in agricultural settings. At this point in time, it is unknown what the effect of the treatment is at the molecular level, but a larger transcriptome study is planned in the near future. This will determine the effect of the treatments on plant genes contributing to colonization, systemic resistance, and plant hormones, as well as genes encoding virulence factors of *P. syringae* pv. *tomato* DC3000.

## Phyllosphere and rhizosphere

To extend the perspective from looking at the experiment as a tri-partite relationship between two bacterial and one plant species to viewing the tomatoes as one system, the effect of the treatments on phyllosphere and rhizosphere was determined. Comparing the microbiome of leaves to that of roots, a first noticeable difference was the extremely low level of 16S rDNA from any bacteria in leaf homogenates from un-challenged tomatoes. Low diversity of microbes in the tomato phyllosphere has been observed before (Dong et al. 2019) and could be due to a masking of bacterial 16S rDNA by high levels of chloroplasts (Tian et al. 2017), as was seen in our experiment. As a general rule, an abundance of bacteria of 10^6^ in leaf homogenate samples used for 16S rDNA sequencing is considered the lower level of detection (Villette et al. 2021). In root homogenates, 16S rDNA was detected in all test groups. A second large difference between phyllosphere and rhizosphere was richness. 684 starting phyllosphere taxonomies were narrowed down to 67 after applying the ≥50 filter. 8,619 rhizosphere taxonomies were narrowed down to 280, applying the much more restrictive ≥500 filter. As one similarity between the two microbiomes, Proteobacteria were the pre-dominant phylum in both.

In phyllosphere and rhizosphere, the microbial composition in this study was similar to previous reports, which also increased confidence that sterilization of the growth medium at the beginning of plant growth would not interfere with the microbiome several weeks later. This is consistent with a previous study on a prairie grass, where the bacterial communities in the soil were no longer discernable between sterile and non-sterile soil after 7 weeks (Hobbs et al. 2026).

Sterile growth medium was also used in one of the three previous studies that the experimental protocol for the phytopathogen experiment in this study was built upon (Bashan and De-Bashan 2002). In full agreement with the literature on the tomato microbiome, proteobacteria were the predominant phylum (Dong et al. 2019), a majority of the species belong to the Gammaproteobacteria (Romero et al. 2014). Actinobacteriota, Bacteroidota, Verrucomicrobiota, and Patescibacteria were part of the rhizosphere alongside Proteobacteria (Fagnano et al. 2025); Rhizobiales and Myxococcales were also part of the rhizosphere (Cheng et al. 2020). Most Pseudomonadales which scored high abundances in this study were from the genus *Pseudomonas,* which has been previously observed (Dong et al. 2019). *Sphingobium* which scored high abundances in this study has been described as part of the rhizosphere in response to α-tomatin (Takamatsu et al. 2023), where it produces indole 3-acetic acid (Gao et al. 2023), and catalyzes toxic steroidal glycoalkaloids (Nakayasu et al. 2023). In contradiction to the literature (Fagnano et al. 2025), Firmicutes could not be identified.

The key part of the microbiome analysis was to characterize the effect the combination treatments had on phyllosphere and rhizosphere. In summary from the phyllosphere, there are three distinct outcomes of the treatments on specific bacterial orders; first, tomato seedling exudate increased the abundance of Enterobacterales in the presence of *A. brasilense* Sp7.

Second, cytidine increased the abundance of Pseudomonadales, more so in the presence of *A. brasilense* Sp7. Third, there is a possibility that *A. brasilense* Sp7 decreased the abundance of Xanthomonadales in all three comparisons between a -*Abra* and a +*Abra* test group.

Analyzing 16S rDNA abundance within the rhizosphere, absence or presence of *A. brasilense* Sp7 appeared to be the largest driver behind the PC1 component. Comparing data from the heatmap to those from the PCA analysis, there is a shift from Flavobacteriales, Pseudomonadales, Propionibacteriales, Saccharomonadales, Rhizobiales, Tepidisphaerales, and Abditibacteriales in the absence of the PGPR to Micrococcales, Azospirillales, Salinisphaerales, and Microtrichales in the presence of *A. brasilense* Sp7. It is especially intriguing that *A. brasilense* increased the abundance of members of its own order Azospirillales, an observation that has not been made before. Furthermore, this appears to go to the expense of Rhizobiales, another large group of beneficial bacteria, though with a strong host specificity for legumes (Boivin et al. 2021; Sijilmassi et al. 2020). This raises the question whether *A. brasilense* may be the perfect PGPR for tomatoes. The authors look forward to testing different cytidine containing inoculants for commercial purposes.

## Conclusion

In conclusion, the combination treatments of *A. brasilense* Sp7 and exudate or *A. brasilense* Sp7 and cytidine had major impacts on the investigated system consisting of tomato plants and the phytopathogen *P. syringae* pv. *tomato* DC3000. A primary observation is the massive reduction by the treatments in disease severity caused by the pathogen. While there is currently not a good hypothesis on the underlying reasons for this reduction, reduction of disease symptoms appeared to be paralleled by an increase in the abundance of the PGPR in the presence of cytidine, which can cause a general improvement of plant health or more specific effects, which will be investigated in future experiments. There was no reduction in the abundance of the challenge pathogen.

## Acknowledgements

The authors thank Dr. Gladys Alexandre (University of Tennessee, Knoxville, TN) for providing the *A. brasilense* Sp7 strain and Dr. Melania Filiatrault (USDA/ARA, Ithaca, NY) for many helpful discussions. We thank Scott Hoselton and Kaycie Schmidt from the Dr. Thomas Glass Biotech Innovation Core at NDSU for performing the 16S rDNA analysis and helping us with the dPCR. We thank Carl Denton for editing the French version of the abstract after producing it with Copilot.

## Competing Interest Statement

The authors declare there are no competing interests.

## Author Contribution Statement

B.M.P. did conceptualization of the research, funding acquisition, project administration, and data curation. B.M.P. wrote the first draft of the manuscript. S.M.H. was responsible for investigation, methodology, student supervision, and reviewing/editing. J.A.N.T., did investigation, data curation, and reviewing/editing. B.O. contributed to investigation and data analysis, and did reviewing/editing.

## Data Availability Statement

All data are either included in the article or provided as online supplement.

## Funding Statement

The research was funded by Specialty Crop Block Grant 22-242 from the USDA/NIFA and grant 24-2030 from the North Dakota Agricultural Products and Utilization Commission. BMP was supported, in part, by the intramural research program of the U.S. Department of Agriculture, National Institute of Food and Agriculture, Hatch project accession number, 7009493. The findings and conclusions in this preliminary publication have not been formally disseminated by the U. S. Department of Agriculture and should not be construed to represent any agency determination or policy.

## Notes

### Competing Interest Statement

The authors have declared no competing interest.

### Summary of Updates

Can. J. Microbiol. has accpeted the manuscript pending minor revisions. We followed their comments.

## References

1. Alexandre, G. 2026. Chemotaxis and other forms of taxis behaviors. *In* The definitive hHandbook of Azospirillum: 100 Years of research and application around the world. *Edited by* F.D. Cassán and L. de Bashan and F. Wisniewski-Dyé and G. A. Maroniche. Springer Nature Switzerland, Cham. pp. 111–125.

2. Alexandre, G., Greer, S.E., and Zhulin, I.B. 2000. Energy taxis is the dominant behavior in *Azospirillum brasilense*. J. Bacteriol. 182(21): 6042–6048. doi:doi:10.1128/JB.182.21.6042-6048.2000.

3. Bashan, Y., and De-Bashan, L.E. 2002. Protection of tomato seedlings against infection by *Pseudomonas syringae* pv. *tomato* by using the plant growth-promoting bacterium *Azospirillum brasilense*. Appl Environ Microbiol 68(6): 2637–2643. doi:10.1128/aem.68.6.2637-2643.2002.

4. Bergougnoux, V. 2014. The history of tomato: from domestication to biopharming. Biotechnol Adv 32(1): 170–189. doi:10.1016/j.biotechadv.2013.11.003.

5. Beringer, J.E. 1974. R factor transfer in *Rhizobium leguminosarum*. Microbiology 84(1): 188–198. doi:doi: 10.1099/00221287-84-1-188.

6. Boivin, S., Mahé, F., Debellé, F., Pervent, M., Tancelin, M., Tauzin, M., Wielbo, J., Mazurier, S., Young, P., and Lepetit, M. 2021. Genetic variation in host-specific competitiveness of the symbiont *Rhizobium leguminosarum* Symbiovar *viciae*. Front Plant Sci 12: 719987. doi:10.3389/fpls.2021.719987.

7. Botta, A.L., Santacecilia, A., Ercole, C., Cacchio, P., and Del Gallo, M. 2013. *In vitro* and *in vivo* inoculation of four endophytic bacteria on *Lycopersicon esculentum*. N Biotechnol 30(6): 666–674. doi:10.1016/j.nbt.2013.01.001.

8. Buell, C.R., Joardar, V., Lindeberg, M., Selengut, J., Paulsen, I.T., Gwinn, M.L., Dodson, R.J., Deboy, R.T., Durkin, A.S., Kolonay, J.F., Madupu, R., Daugherty, S., Brinkac, L., Beanan, M.J., Haft, D.H., Nelson, W.C., Davidsen, T., Zafar, N., Zhou, L., Liu, J., Yuan, Q., Khouri, H., Fedorova, N., Tran, B., Russell, D., Berry, K., Utterback, T., Van Aken, S.E., Feldblyum, T.V., D’Ascenzo, M., Deng, W.L., Ramos, A.R., Alfano, J.R., Cartinhour, S., Chatterjee, A.K., Delaney, T.P., Lazarowitz, S.G., Martin, G.B., Schneider, D.J., Tang, X., Bender, C.L., White, O., Fraser, C.M., and Collmer, A. 2003. The complete genome sequence of the *Arabidopsis* and tomato pathogen *Pseudomonas syringae* pv. *tomato* DC3000. Proc Natl Acad Sci U S A 100(18): 10181–10186. doi:10.1073/pnas.1731982100.

9. Cassán, F., Vanderleyden, J., and Spaepen, S. 2014. Physiological and agronomical aspects of phytohormone production by model plant-growth-promoting rhizobacteria (PGPR) belonging to the genus *Azospirillum*. J Plant Growth Regul 33(2): 440–459.

10. Cassán, F., Maroniche, G., Reis, V.M., Etto, R.M., Bellotti, G., Alexandre, G., Torres, D., Nievas, S., Tripathi, A.K., Perez, R., de Souza, E.M., Pineda, E.G., Pedraza, R., Peña, E.M.B., Gureeva, M., Prigent-Combaret, C., Díaz-Zorita, M., Vio, S.A., Del Gallo, M., Buttrós, V.H., de Oliveira Pedrosa, F., Puente, M., García, J.E., López, B., Palacios, O., Wisniewski-Dyé, F., and de-Bashan, L. 2026. A century of research on *Azospirillum* and still so much to discover. Plant and Soil. doi:10.1007/s11104-025-08178-9.

11. Cassán, F.D.d.B., L.; Wisniewski-Dyé, F.; Maroniche, G.A. 2026. The definitive handbook of Azospirillum. Springer.

12. Cerna-Vargas, J.P., Santamaría-Hernando, S., Matilla, M.A., Rodríguez-Herva, J.J., Daddaoua, A., Rodríguez-Palenzuela, P., Krell, T., and López-Solanilla, E. 2019. Chemoperception of specific amino acids controls phytopathogenicity in *Pseudomonas syringae* pv. tomato. mBio 10(5). doi:10.1128/mBio.01868-19.

13. Chakravarthy, S., Butcher, B.G., Liu, Y., D’Amico, K., Coster, M., and Filiatrault, M.J. 2017. Virulence of *Pseudomonas syringae* pv. tomato DC3000 Is influenced by the catabolite repression control protein Crc. Mol Plant Microbe Interact 30(4): 283–294. doi:10.1094/mpmi-09-16-0196-r.

14. Cheng, Z., Lei, S., Li, Y., Huang, W., Ma, R., Xiong, J., Zhang, T., Jin, L., Haq, H.U., Xu, X., and Tian, B. 2020. Revealing the variation and stability of bacterial communities in tomato rhizosphere microbiota. Microorganisms 8(2). doi:10.3390/microorganisms8020170.

15. Compant, S., Cassan, F., Kostić, T., Johnson, L., Brader, G., Trognitz, F., and Sessitsch, A. 2025. Harnessing the plant microbiome for sustainable crop production. Nature Rev Microbiol 23(1): 9–23. doi:10.1038/s41579-024-01079-1.

16. Coniglio, A., Larama, G., Molina, R., Mora, V., Torres, D., Marin, A., Avila, A.I., Lede NoirCarlan, C., Erijman, L., and Figuerola, E.L. 2022. Modulation of maize rhizosphere microbiota composition by inoculation with *Azospirillum argentinense* Az39 (formerly *A. brasilense* Az39). J Soil Sci Plant Nutr 22(3): 3553–3567.

17. Contreras-Cornejo, H.A., Sanchez-Yanez, J.M.; Carrillo-Flores, E.; Mellado-Rojas, Ma.E.; Berltran-Pena, E.M. 2026. Interaction of Azospirillum ssp. with plants; molecular plant growth responses. *In* The definitive hHandbook of Azospirillum: 100 Years of research and application around the world. *Edited by* F.D. Cassán and L. de Bashan and F. Wisniewski-Dyé and G. A. Maroniche. Springer Nature Switzerland, Cham. pp. 323–342.

18. da Cunha, E.T., Pedrolo, A.M., Paludo, F., Scariot, M.C., and Arisi, A.C.M. 2020. *Azospirillum brasilense* viable cells enumeration using propidium monoazide-quantitative PCR. Arch Microbiol 202(7): 1653–1662. doi:10.1007/s00203-020-01877-0.

19. Dong, C.-J., Wang, L.-L., Li, Q., and Shang, Q.-M. 2019. Bacterial communities in the rhizosphere, phyllosphere and endosphere of tomato plants. PloS one 14(11): e0223847.

20. El-Beltagi, H.S., Ahmad, I., Basit, A., Abd El-Lateef, H.M., Yasir, M., Tanveer Shah, S., Ullah, I., Elsayed Mohamed Mohamed, M., Ali, I., Ali, F., Ali, S., Aziz, I., Kandeel, M., and Zohaib Ikram, M. 2022. Effect of *Azospirillum* and *Azotobacter* species on the performance of cherry tomato under different salinity levels. Gesunde Pflanzen 74(2): 487–499. doi:doi:10.1007/s10343-022-00625-2.

21. El-Fatah, B., Imran, M., Abo-Elyousr, K.A.M., and Mahmoud, A.F. 2023. Isolation of *Pseudomonas syringae* pv. tomato strains causing bacterial speck disease of tomato and marker-based monitoring for their virulence. Mol Biol Rep 50(6): 4917–4930. doi:10.1007/s11033-023-08302-x.

22. Elías, J.M., Ramírez-Mata, A., Albornóz, P.L., Baca, B.E., Díaz-Ricci, J.C., and Pedraza, R.O. 2022. The polar flagellin of *Azospirillum brasilense* REC3 induces a defense response in strawberry plants against the fungus *Macrophomina phaseolina*. J Plant Gro Regul 41(7): 2992–3008. doi:10.1007/s00344-021-10490-4.

23. Elsharkawy, M.M., Khedr, A.A., Mehiar, F., El-Kady, E.M., Alwutayd, K.M., and Behiry, S.I. 2023. Rhizobacterial colonization and management of bacterial speck pathogen in tomato by *Pseudomonas* spp. Microorganisms 11(5). doi:10.3390/microorganisms11051103.

24. Eskew, D.F., D.D. 1977. Nitrogen fixation, denitrification, and pleomorphic growth in a highly pigmented *Spirillum lipoferum*. Appl. Environ. Microbiol. 34: 582–585. doi:doi: 10.1128/aem.34.5.582-585.1977.

25. Fagnano, F.M., Ventorino, V., Pasolli, E., Romano, I., Ambrosino, P., and Pepe, O. 2025. From microbiome to biostimulants: unlocking the potential of tomato root endophytes. BMC Plant Biol 25(1): 427. doi:10.1186/s12870-025-06447-4.

26. Feng, Z., Liang, Q., Yao, Q., Bai, Y., and Zhu, H. 2024. The role of the rhizobiome recruited by root exudates in plant disease resistance: current status and future directions. Environ Microbiome 19(1): 91. doi:10.1186/s40793-024-00638-6.

27. Fukami, J., de la Osa, C., Ollero, F.J., Megías, M., and Hungria, M. 2018. Co-inoculation of maize with *Azospirillum brasilense* and *Rhizobium tropici* as a strategy to mitigate salinity stress. Funct Plant Biol 45(3): 328–339. doi:10.1071/fp17167.

28. Ganusova, E.E., Russell, M.H., Patel, S., Seats, T., and Alexandre, G. 2024. An *Azospirillum brasilense* chemoreceptor that mediates nitrate chemotaxis has conditional roles in the colonization of plant roots. Appl Environ Microbiol 90(6): e0076024. doi:10.1128/aem.00760-24.

29. Ganusova, E.E., Vo, L.T., Abraham, P.E., O’Neal Yoder, L., Hettich, R.L., and Alexandre, G. 2021. The *Azospirillum brasilense* core chemotaxis proteins CheA1 and CheA4 link chemotaxis signaling with nitrogen metabolism. mSystems 6(1). doi:10.1128/mSystems.01354-20.

30. Ganusova, E.E., Rost, M., Aksenova, A., Abdulhussein, M., Holden, A., and Alexandre, G. 2023. *Azospirillum brasilense* AerC and Tlp4b cytoplasmic chemoreceptors are promiscuous and interact with the two membrane-bound chemotaxis signaling clusters mediating chemotaxis responses. J Bacteriol 205(6): e0048422. doi:10.1128/jb.00484-22.

31. Gao, R., Dong, H., Liu, Y., Yao, Q., Li, H., and Zhu, H. 2023. *Sphingomonas lycopersici* sp. *nov*., isolated from tomato rhizosphere soil. Int J Syst Evol Microbiol 73(5). doi:10.1099/ijsem.0.005920.

32. Hobbs, A., Ochoa-Rojas, D., Humphrey, C.E., Kyndt, J.A., and Moore, T.C. 2026. Soil microbiome perturbation impedes growth of Bouteloua curtipendula and increases relative abundance of soil microbial pathogens. PLoS One 21(1): e0312218. doi:10.1371/journal.pone.0312218.

33. Housh, A.B., Noel, R., Powell, A., Waller, S., Wilder, S.L., Sopko, S., Benoit, M., Powell, G., Schueller, M.J., and Ferrieri, R.A. 2023. Studies Using Mutant Strains of Azospirillum brasilense Reveal That Atmospheric Nitrogen Fixation and Auxin Production Are Light Dependent Processes. Microorganisms 11(7): 1727. Available from https://www.mdpi.com/2076-2607/11/7/1727 [accessed.

34. Huang, J., Zhu, L., Lu, X., Cui, F., Wang, J., and Zhou, C. 2023. A simplified synthetic rhizosphere bacterial community steers plant oxylipin pathways for preventing foliar phytopathogens. Plant Physiol Biochem 202: 107941. doi:10.1016/j.plaphy.2023.107941.

35. Janda, M., Rybak, K., Krassini, L., Meng, C., Feitosa-Junior, O., Stigliano, E., Szulc, B., Sklenar, J., Menke, F.L.H., Malone, J.G., Brachmann, A., Klingl, A., Ludwig, C., and Robatzek, S. 2023. Biophysical and proteomic analyses of *Pseudomonas syringae* pv. *tomato* DC3000 extracellular vesicles suggest adaptive functions during plant infection. mBio 14(4): e0358922. doi:10.1128/mbio.03589-22.

36. Johnsen, K., and Nielsen, P. 1999. Diversity of *Pseudomonas* strains isolated with King’s B and Gould’s S1 agar determined by repetitive extragenic palindromic-polymerase chain reaction, 16S rDNA sequencing and Fourier transform infrared spectroscopy characterisation. FEMS Microbiol Lett 173(1): 155–162. doi:10.1111/j.1574-6968.1999.tb13497.x.

37. Keren, G., Yehezkel, G., Satish, L., Adamov, Z., Barak, Z.e., Ben-Shabat, S., Kagan-Zur, V., and Sitrit, Y. 2024. Root-secreted nucleosides: signaling chemoattractants of rhizosphere bacteria [Original Research]. Frontiers in Plant Science Volume 15 - 2024. doi:10.3389/fpls.2024.1388384.

38. King, E.O., Ward, M.K., and Raney, D.E. 1954. Two simple media for the demonstration of pyocyanin and fluorescin. J Lab Clin Med 44(2): 301–307.

39. King, W.L., Grandinette, E.M., Trase, O., Rolon, M.L., Salis, H.M., Wood, H., and Bell, T.H. 2024. Autoclaving is at least as effective as gamma irradiation for biotic clearing and intentional microbial recolonization of soil. mSphere 9(7): e00476–00424. doi:doi:10.1128/msphere.00476-24.

40. Korenblum, E., Dong, Y., Szymanski, J., Panda, S., Jozwiak, A., Massalha, H., Meir, S., Rogachev, I., and Aharoni, A. 2020. Rhizosphere microbiome mediates systemic root metabolite exudation by root-to-root signaling. Proc Natl Acad Sci U S A 117(7): 3874–3883. doi:10.1073/pnas.1912130117.

41. Lindeberg, M., Biehl, B.S., Glasner, J.D., Perna, N.T., Collmer, A., and Collmer, C.W. 2009. Gene ontology annotation highlights shared and divergent pathogenic strategies of type III effector proteins deployed by the plant pathogen *Pseudomonas syringae* pv *tomato* DC3000 and animal pathogenic *Escherichia coli strains*. BMC Microbiol 9 Suppl 1(Suppl 1): S4. doi:10.1186/1471-2180-9-s1-s4.

42. Ma, L.J., Geiser, D.M., Proctor, R.H., Rooney, A.P., O’Donnell, K., Trail, F., Gardiner, D.M., Manners, J.M., and Kazan, K. 2013. *Fusarium* pathogenomics. Annu Rev Microbiol 67: 399–416. doi:10.1146/annurev-micro-092412-155650.

43. Ma, S., Chen, Q., Zheng, Y., Ren, T., He, R., Cheng, L., Zou, P., Jing, C., Zhang, C., and Li, Y. 2025. A tale for two roles: Root-secreted methyl ferulate inhibits *P. nicotianae* and enriches the rhizosphere *Bacillus* against black shank disease in tobacco. Microbiome 13(1): 33. doi:10.1186/s40168-024-02008-3.

44. Mora, V., López, G., Molina, R., Coniglio, A., Nievas, S., De Diego, N., Zeljković, S.Ć., Sarmiento, S.S., Spíchal, L., Robertson, S., Wilkins, O., Elías, J., Pedraza, R., Estevez, J.M., Belmonte, M.F., and Cassán, F. 2023. *Azospirillum argentinense* modifies *Arabidopsis* root architecture through auxin-dependent pathway and flagellin. Journal of Soil Science and Plant Nutrition 23(3): 4543–4557. doi:10.1007/s42729-023-01371-8.

45. Nakayasu, M., Takamatsu, K., Yazaki, K., and Sugiyama, A. 2022. Plant specialized metabolites in the rhizosphere of tomatoes: secretion and effects on microorganisms. Biosci Biotechnol Biochem 87(1): 13–20. doi:10.1093/bbb/zbac181.

46. Nakayasu, M., Ohno, K., Takamatsu, K., Aoki, Y., Yamazaki, S., Takase, H., Shoji, T., Yazaki, K., and Sugiyama, A. 2021. Tomato roots secrete tomatine to modulate the bacterial assemblage of the rhizosphere. Plant Physiol 186(1): 270–284. doi:10.1093/plphys/kiab069.

47. Nakayasu, M., Takamatsu, K., Kanai, K., Masuda, S., Yamazaki, S., Aoki, Y., Shibata, A., Suda, W., Shirasu, K., Yazaki, K., and Sugiyama, A. 2023. Tomato root-associated *Sphingobium* harbors genes for catabolizing toxic steroidal glycoalkaloids. mBio 14(5): e0059923. doi:10.1128/mbio.00599-23.

48. Namgung, M., Lim, Y.J., Kang, M.K., Oh, C.S., and Park, D.H. 2019. *Pseudomonas syringae* pv. *tomato* DC3000 improves *Escherichia coli* O157:H7 survival in tomato plants. J Microbiol Biotechnol 29(12): 1975–1981. doi:10.4014/jmb.1907.07072.

49. Nievas, S., Coniglio, A., Takahashi, W.Y., López, G.A., Larama, G., Torres, D., Rosas, S., Etto, R.M., Galvão, C.W., Mora, V., and Cassán, F. 2023. Unraveling *Azospirillum*’s colonization ability through microbiological and molecular evidence. J Appl Microbiol 134(4). doi:10.1093/jambio/lxad071.

50. Nisha, F.A., Horne, S.M., and Prüß, B.M. 2025. *Azospirillum brasilense* and cytidine enhance lateral roots of peas. FEMS Microbiol Lett 372: fnaf025. doi:10.1093/femsle/fnaf025.

51. Nisha, F.A., Tagoe, J.N.A., Pease, A.B., Horne, S.M., Ugrinov, A., Geddes, B.A., and Prüß, B.M. 2024. Plant seedlings of peas, tomatoes, and cucumbers exude compounds that are needed for growth and chemoattraction of *Rhizobium leguminosarum* bv. *viciae* 3841 and *Azospirillum brasilense* Sp7. Can J Microbiol 70(5): 150–162. doi:10.1139/cjm-2023-0217.

52. Pellegrini, M., Ercole, C., Di Zio, C., Matteucci, F., Pace, L., and Del Gallo, M. 2020. *In vitro* and *in planta* antagonistic effects of plant growth-promoting rhizobacteria consortium against soilborne plant pathogens of *Solanum tuberosum* and *Solanum lycopersicum*. FEMS Microbiol. Lett. 367(13). doi:doi:10.1093/femsle/fnaa099.

53. Peňázová, E., Dvořák, M., Ragasová, L., Kiss, T., Pečenka, J., Čechová, J., and Eichmeier, A. 2020. Multiplex real-time PCR for the detection of *Clavibacter michiganensis* subsp. *michiganensis*, *Pseudomonas syringae* pv. *tomato* and pathogenic *Xanthomonas* species on tomato plants. PLoS One 15(1): e0227559. doi:10.1371/journal.pone.0227559.

54. Pereg-Gerk, L., Paquelin, A., Gounon, P., Kennedy, I.R., and Elmerich, C. 1998. A transcriptional regulator of the LuxR-UhpA family, FlcA, controls flocculation and wheat root surface colonization by *Azospirillum brasilense* Sp7. Mol Plant Microbe Interact 11(3): 177–187. doi:10.1094/mpmi.1998.11.3.177.

55. Querejeta, G.A. 2023. Sterilize methods comparison for soils: cost, time, and efficiency. Internat J Methodol 2(1): 34–40. doi:10.21467/ijm.2.1.6263.

56. Reddy Priya, P., Selastin Antony, R., Gopalaswamy, G., and Balachandar, D. 2016. Development of sequence-characterized amplified region (SCAR) markers as a quality standard of inoculants based on *Azospirillum*. Arch Microbiol 198(3): 257–267. doi:10.1007/s00203-016-1187-7.

57. Romero, F.M., Marina, M., and Pieckenstain, F.L. 2014. The communities of tomato (*Solanum lycopersicum L*.) leaf endophytic bacteria, analyzed by 16S-ribosomal RNA gene pyrosequencing. FEMS Microbiol Lett 351(2): 187–194. doi:10.1111/1574-6968.12377.

58. Sijilmassi, B., Filali-Maltouf, A., Boulahyaoui, H., Kricha, A., Boubekri, K., Udupa, S., Kumar, S., and Amri, A. 2020. Assessment of genetic diversity and symbiotic efficiency of selected Rhizobia strains nodulating lentil (*Lens culinaris* Medik.). Plants (Basel) 10(1). doi:10.3390/plants10010015.

59. Steenhoudt, O., and Vanderleyden, J. 2000. *Azospirillum*, a free-living nitrogen-fixing bacterium closely associated with grasses: genetic, biochemical and ecological aspects. FEMS Microbiol Rev 24(4): 487–506.

60. Takamatsu, K., Toyofuku, M., Okutani, F., Yamazaki, S., Nakayasu, M., Aoki, Y., Kobayashi, M., Ifuku, K., Yazaki, K., and Sugiyama, A. 2023. α-Tomatine gradient across artificial roots recreates the recruitment of tomato root-associated *Sphingobium*. Plant Direct 7(12): e550. doi:10.1002/pld3.550.

61. Tian, X., Shi, Y., Geng, L., Chu, H., Zhang, J., Song, F., Duan, J., and Shu, C. 2017. Template preparation affects 16S rRNA high-throughput sequencing analysis of phyllosphere microbial communities. Front Plant Sci 8: 1623.

62. Vazquez, A., Zawoznik, M., Benavides, M.P., and Groppa, M.D. 2021. *Azospirillum brasilense* Az39 restricts cadmium entrance into wheat plants and mitigates cadmium stress. Plant Sci. 312: 111056. doi:doi:10.1016/j.plantsci.2021.111056.

63. Vidotti, M.S., Matias, F.I., Alves, F.C., Pérez-Rodríguez, P., Beltran, G.A., Burgueño, J., Crossa, J., and Fritsche-Neto, R. 2019. Maize responsiveness to *Azospirillum brasilense*: Insights into genetic control, heterosis and genomic prediction. PLoS One 14(6): e0217571. doi:doi:10.1371/journal.pone.0217571.

64. Villette, R., Autaa, G., Hind, S., Holm, J.B., Moreno-Sabater, A., and Larsen, M. 2021. Refinement of 16S rRNA gene analysis for low biomass biospecimens. Sci Rep 11(1): 10741. doi:10.1038/s41598-021-90226-2.

65. Wang, H., Jin, H., Chen, Z., Li, W., Ma, J., Hu, T., Liu, Q., Zhang, Y., Lin, X., and Xie, Z. 2024. *Azospirillum isscasi* sp. nov., a bacterium isolated from rhizosphere soil of rice. Int J Syst Evol Microbiol 74(1). doi:10.1099/ijsem.0.006218.

66. Wei, H.L., and Collmer, A. 2018. Defining essential processes in plant pathogenesis with *Pseudomonas syringae* pv. *tomato* DC3000 disarmed polymutants and a subset of key type III effectors. Mol Plant Pathol 19(7): 1779–1794. doi:10.1111/mpp.12655.

67. Whalen, M.C., Innes, R.W., Bent, A.F., and Staskawicz, B.J. 1991. Identification of *Pseudomonas syringae* pathogens of *Arabidopsi*s and a bacterial locus determining avirulence on both *Arabidopsis* and soybean. Plant Cell 3(1): 49–59. doi:10.1105/tpc.3.1.49.

68. Xin, X.F., and He, S.Y. 2013. *Pseudomonas syringae* pv. tomato DC3000: a model pathogen for probing disease susceptibility and hormone signaling in plants. Annu Rev Phytopathol 51: 473–498. doi:10.1146/annurev-phyto-082712-102321.

